# The molecular pathogenesis of superoxide dismutase 1-linked ALS is promoted by low oxygen tension

**DOI:** 10.1101/320283

**Authors:** Isil Keskin, Elin Forsgren, Manuela Lehmann, Peter M. Andersen, Thomas Brännström, Dale J. Lange, Matthis Synofzik, Ulrika Nordström, Per Zetterström, Stefan L. Marklund, Jonathan D. Gilthorpe

## Abstract

Mutations that destabilize superoxide dismutase 1 (SOD1) are a cause of amyotrophic lateral sclerosis (ALS). SOD1, which is located in the reducing cytosol, contains an oxidized disulfide bond required for stability. We show that the bond is an Achilles heel of the protein because it is sensitive to the oxygen tension. Culture of ALS patient-derived fibroblasts, astrocytes and induced pluripotent stem cell-derived mixed motor neuron and astrocyte cultures (MNACs) under lowered oxygen tensions caused reductive bond cleavage and misfolding. The effects were greatest in cells expressing mutant SOD1s, but also occurred in wild type SOD1 in cultures derived from patients carrying ALS-linked mutations in *C9orf72, FUS* and *TBK1*, as well as from controls. MNACs showed a greater response than the other cell types, including enhanced SOD1 aggregation, in line with the vulnerability of the motor system. Our results show that oxygen tension is a principal determinant of SOD1 stability and shed light on how risk factors for ALS, such as aging and other conditions causing reduced vascular perfusion, could lead to disease initiation and progression.

**Subject categories** Neuroscience; Molecular Biology of Disease

## Introduction

Amyotrophic lateral sclerosis (ALS) is characterized by adult-onset progressive degeneration of upper and lower motor neurons (MN) and their associated tracts. Usually the disease begins focally and then spreads contiguously, resulting in progressive paralysis and death from respiratory failure (Charcot, 1873). Mutations in the gene encoding the ubiquitously expressed, free radical scavenging enzyme superoxide dismutase 1 (SOD1) cause ALS (Rosen et al., 1993), and are found in 1-9% of patients (Andersen & Al-Chalabi, 2011). Of over 200 mutations identified, 174 are missense and result in SOD1 variants of which most in large, although variable proportions are natively folded (Andersen et al., 1995, Sato et al., 2005, Synofzik et al., 2012). Twenty-seven encode mutants with disruptive changes (insertions, deletions and truncations) that impede native folding. Most of the truncated mutants lack C146 and thereby the C57-C146 disulfide bond, which is critical for SOD1 stability. These mutants are highly unstable and rapidly degraded (Birve et al., 2010, Jonsson et al., 2004, Keskin et al., 2017, Keskin et al., 2016, Sato et al., 2005). Hence, the collective genetic and biochemical evidence from patients infers that any common ALS-causing SOD1 species is disordered, lacks the C57-C146 disulfide bond and is only present in minute amounts.

Neuronal inclusions containing aggregated SOD1 are a hallmark of ALS, both in patients and in transgenic (Tg) animal models expressing mutant human SOD1s (Kato et al., 2000). We have shown that two structurally distinct strains of SOD1 aggregates form in Tg mouse models (Bergh et al., 2015), and that both transmit spreading, templated SOD1 aggregation and fatal ALS-like disease in a prion-like manner (Bidhendi et al., 2016). Similarly, others have shown seeding effects of spinal cord extracts from end-stage SOD1 Tg mice (Ayers et al., 2014, Ayers et al., 2016). The templated growth of SOD1 aggregation depends on disordered, disulfide-reduced SOD1 species as building blocks (Chattopadhyay et al., 2008, Chattopadhyay et al., 2015, Furukawa et al., 2008, Lang et al., 2012). This could represent the core pathogenic mechanism in SOD1-induced ALS.

Human SOD1 is primarily localized in the cytosol and is composed of two equal 153 amino acid-long subunits, each containing 1 Cu and 1 Zn ion as well as the stabilizing C57-C146 disulfide bond. SOD1 is an ancient enzyme and was secreted to the oxidizing extracellular/periplasmic space in early unicellular organisms, as well as in current α-bacteria (Bordo et al., 1994, Miller, 2012). However, the cytosol is strongly reducing and the C57-C146 bond could be an evolution-related Achilles heel of eukaryotic SOD1, and a critical determinant of its role in ALS. Absence of the disulfide bond promotes misfolding and aggregation of SOD1 *in vitro* (Chattopadhyay et al., 2008, Chattopadhyay et al., 2015, Furukawa et al., 2008, Keskin et al., 2016, Lang et al., 2012). Moreover, aggregated as well as soluble misfolded SOD1 species in the central nervous system (CNS) of SOD1 Tg models lack the bond (Bergemalm et al., 2010, Karch et al., 2009, Zetterström et al., 2013, Zetterstrom et al., 2007).

If the C57-C146 disulfide bond in SOD1 were in equilibrium with reduced and oxidized glutathione (GSH/GSSG) -the principal redox couple in the cytosol - it would be almost completely reduced (Mercatelli et al., 2016, Schwarzlander et al., 2016). Instead, the status of C57-C146 seems to be determined kinetically with oxidation provided by O_2_ and catalyzed by copper chaperone for SOD (CCS) (Culotta et al., 1997, Fetherolf et al., 2017, Furukawa et al., 2004). Reduction is provided by glutaredoxin-1 using reducing equivalents from GSH (Carroll et al., 2006). Thus, cellular O_2_ tension might be a major determinant of SOD1 disulfide bond stability.

Age is the principal non-hereditary risk factor for ALS, as well as most other neurodegenerative diseases. It is associated with reduced perfusion in the CNS owing to vascular wall degeneration and neurovascular unit dysfunction (Iadecola, 2017). We hypothesized that local reductions in O_2_ tension caused by impaired perfusion may have an effect on SOD1 stability. To test this idea we used *in vitro* cell culture models of ALS including fibroblasts from ALS patients carrying mutations in *SOD1*, other ALS-linked genes and from non-disease controls. We also examined neural cells including primary patient-derived astrocytes and mixed motor neuron and astrocyte cultures (MNACs) derived from induced pluripotent stem cells (iPSCs). We found that low O_2_ tensions markedly increased disulfide bond reduction, misfolding and aggregation of SOD1 in a time-and concentration-dependent manner.

## Results

### Low oxygen tensions promote misfolding of SOD1 in patient-derived cells

Previously, we have utilized ALS patient-derived dermal fibroblasts as an *in vitro* model to study misfolding under physiological SOD1 expression levels (Keskin et al., 2016). Cells cultured *in vitro* are typically maintained in humidified ambient air supplemented with 5% (v/v) CO_2_, resulting in an O_2_ tension of approximately 19%. However, under normal conditions the O_2_ tension in the CNS is much lower, ranging between 0.2 and 5% (Erecinska & Silver, 2001, Lyons et al., 2016). Moreover, an age-related disruption in perfusion might lead to further reductions in local O_2_ tensions. Therefore, we examined the SOD1 misfolding response in fibroblast lines carrying ALS-linked *SOD1* mutations and cultured at reduced O_2_ tensions for 24 h (Fig 1A and Table EV1).

**Figure 1.** Experimental overview and generation of mixed motor neuron and astrocyte cultures (MNACs) derived from induced pluripotent stem cells (iPSCs). (A) Patient-derived fibroblasts, primary spinal cord ventral horn astrocytes and iPSC-derived MNACs were exposed to different O_2_ tensions for 24 h prior to analysis. (B) Patient-derived fibroblasts were reprogrammed to iPSCs using the 4 Yamanaka factors and then differentiated to a forelimb level, ventral spinal cord identity over 14 days. At Day 14, differentiated cells were dissociated and plated on poly-L-ornithine/laminin and matured for 10 more days. MNACs were used for O_2_ tension experiments at Day 25.

Misfolded SOD1 was quantified with a specific ELISA (misELISA) (Zetterström et al., 2011). Lines carrying the *SOD1^N86S^* and *SOD1^E78_R79insSI^* mutations showed distinct O_2_ concentration responses, with the highest levels of misfolding detected at 1% O_2_ (Fig 2A). Interestingly, no increase in SOD1 misfolding was seen in a line carrying the *SOD1^G127Gfs*7^* (G127X) truncating mutation, which results in a protein that lacks the C57-C146 disulfide bond and cannot fold properly (Fig 2A and Fig EV1A).

**Figure 2.**
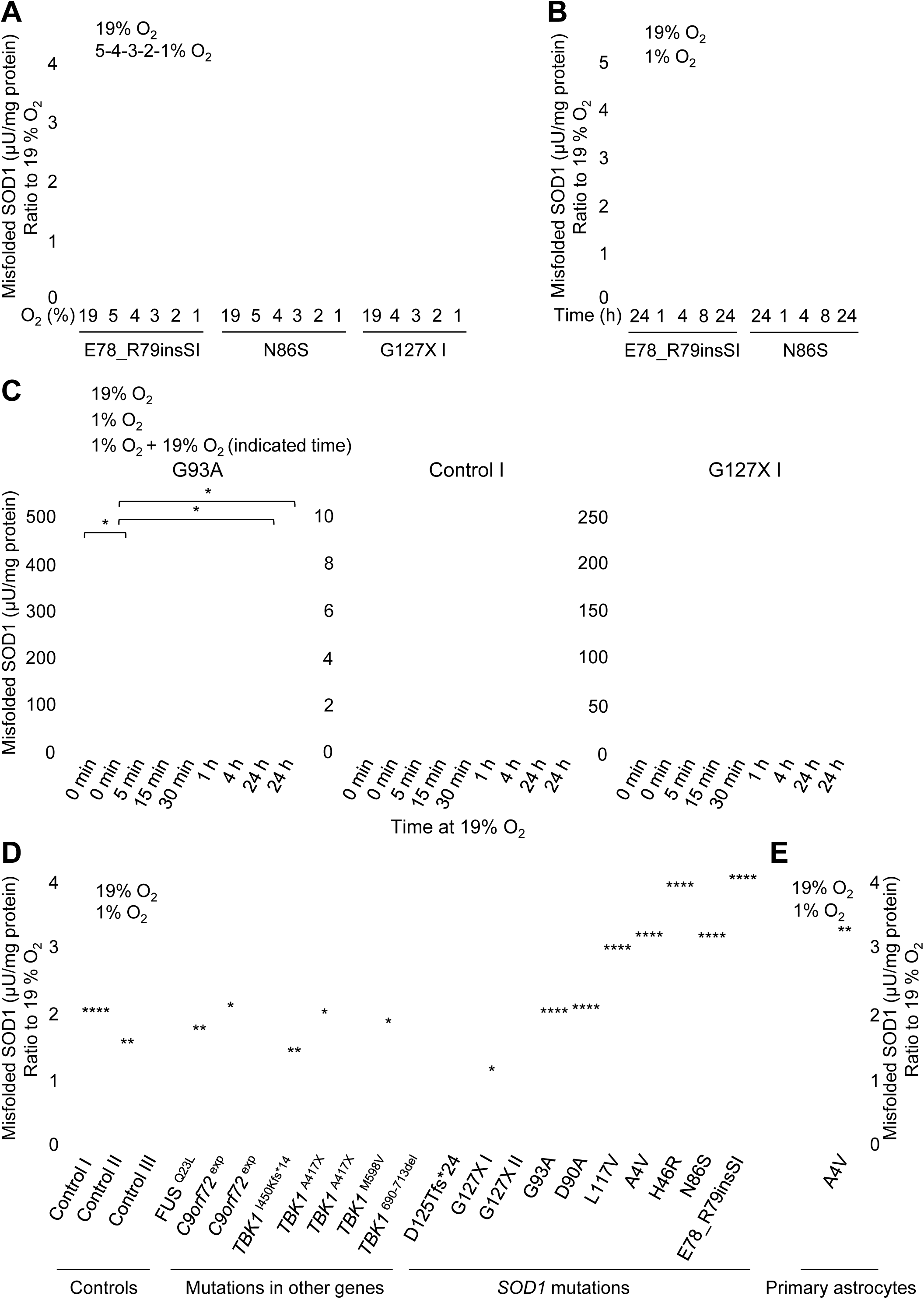
Concentration-and time-dependent misfolding of SOD1 in response to low oxygen tensions. Misfolded SOD1 quantified in fibroblast extracts by misELISA, normalized to total protein and presented as a ratio to the level present in replicate cultures maintained at 19% O_2_ for 24 h. (A) O_2_ concentration- (19-1%) and (B) time- (1-24 h) dependent increases in misfolded SOD1 in fibroblast lines. Data are expressed as the mean ± SD (n = 3 replicate culture wells). (C) Reversal of the misfolding response following transfer of cultures from 1% O_2_/24 h to 19% O_2_. Data are expressed as the mean ± SD (n = 3 replicate culture wells for Control and SOD1^G127X^, 2 replicate experiments each with 3 replicate cultures for G93A), \**p* < 0.05 analyzed by the Kruskal-Wallis test followed by Dunn’s test. (D) Misfolding response in patient-derived fibroblast lines and (E) primary astrocytes following exposure to 1% O_2_ for 24 h. Data are expressed as the mean ± SD (n = 3 to 15; 1-5 replicate experiments each with 3 replicate cultures), \**p* < 0.05, \*\**p* < 0.01, \*\*\*\**p* < 0.0001 analyzed by Mann-Whitney U test. Blue bars = fibroblasts, green bars = astrocytes, grey bars = corresponding cell lines cultured at 19% O_2_.

We next investigated the temporal response of misfolding to low O_2_ and detected a significant increase in misfolded SOD1 at 1% O_2_ after 4 h, reaching a maximum after 8 h, which was maintained at 24 h (Fig 2B). Quantification of total SOD1 revealed that increased misfolding was not due to increased SOD1 expression (Fig EV1B). To address whether the increase in misfolding was reversible, we quantified misfolded SOD1 at different time points after returning cells from 24 h at 1% O_2_ to 19% O_2_. A rapid reversal was observed in the SOD1^G93A^ line with a misfolded SOD1 half-life of ~15 min, returning to baseline after ~1 h (Fig 2C). Reversal of misfolding of wild-type SOD1 (SOD1^WT^) in a control line was even more rapid (Fig 2C). No significant effects were seen in the SOD1^G127X^ line (Fig 2C). Hence, we concluded that the misfolding response was inversely proportional to the O_2_ tension. It was also time-dependent, reversible, and appeared to require the C57-C146 disulfide bond.

To probe the misfolding response in more detail, we next analyzed a panel of control and patient-derived fibroblast lines cultured under conditions where the misfolding response was maximal (1% O_2_, 24 h; Fig 2D and Fig EV1C). A significant increase in misfolded SOD1 was seen in all lines that carried full-length mutant SOD1s and ranged from 1.5 – 3.5-fold. Significant increases of a smaller magnitude were also observed in most control lines, as well as those from patients carrying ALS-linked *C9orf72*, *FUS* or *TBK1* mutations, which only contain SOD1^WT^. The only fibroblasts that did not show significant increases were lines carrying *SOD1* mutations resulting in truncations and the absence of C146*: SOD1^G127X^* and *SOD1^D125Tfs*24^*.

To investigate the misfolding response in patient-derived neural cells, we generated primary astrocytes from the lateral ventral horn of an ALS patient heterozygous for the *SOD1^A4V^* mutation (Fig 1A, Fig 2E, Fig EV1 D and E and Table EV1) (Haidet-Phillips et al., 2011). The level of misfolded SOD1 increased 2-fold when these cells were cultured at 1% O_2_ compared to 19% for 24 h (Fig 2E and Fig EV1D), which was similar to the increase seen in fibroblasts from another *SOD1^A4V^* patient (Fig 2D and Fig EV1C).

### An enhanced SOD1 misfolding response in patient-derived motor neuron/astrocyte cultures

Since ALS preferentially targets the motor system, we reprogrammed a subset of the patient-derived fibroblast lines to iPSCs and differentiated these to MNACs (Fig 1A and B and Table EV1). Recently we have found that SOD1 misfolding is enhanced in MNACs compared to the corresponding fibroblasts, iPSCs and iPSC-derived sensory neurons (Forsgren et al., under revision). After 10 days in culture post-differentiation (Day 25; Fig 1B), MNACs contained ~5% neurons expressing tubulin beta 3 class III (TUBB3; Fig EV2 A and B). A large proportion of these neurons (78 – 84%) co-expressed the motor neuron markers ISL LIM homeobox 1/2 (ISL1/2) and non-phosphorylated neurofilament-H (SMI32), representing a limb innervating subtype that are highly susceptible to ALS (Amoroso et al., 2013).

We first titrated the dose-response of SOD1 misfolding to O_2_ tension in MNACs from control, SOD1^A4V^ and SOD1^G93A^ lines. A significant increase in misfolding was detected at ≤4% and was maximal at 2% O_2_, which was enhanced in MNACs compared to fibroblasts in all three lines (Fig 3A). Under the same conditions used for fibroblasts (1% O_2_ for 24 h), a significant degree of axonal fragmentation was observed (data not shown), indicative of neuronal stress. However, since axonal morphology (Fig EV2B), cellular ATP levels (Fig EV3A) and viability (Fig EV3B) were not significantly affected by exposure to 2% O_2_ for 24 h, this was used to investigate the misfolding response in MNACs from ALS patients.

**Figure 3.**
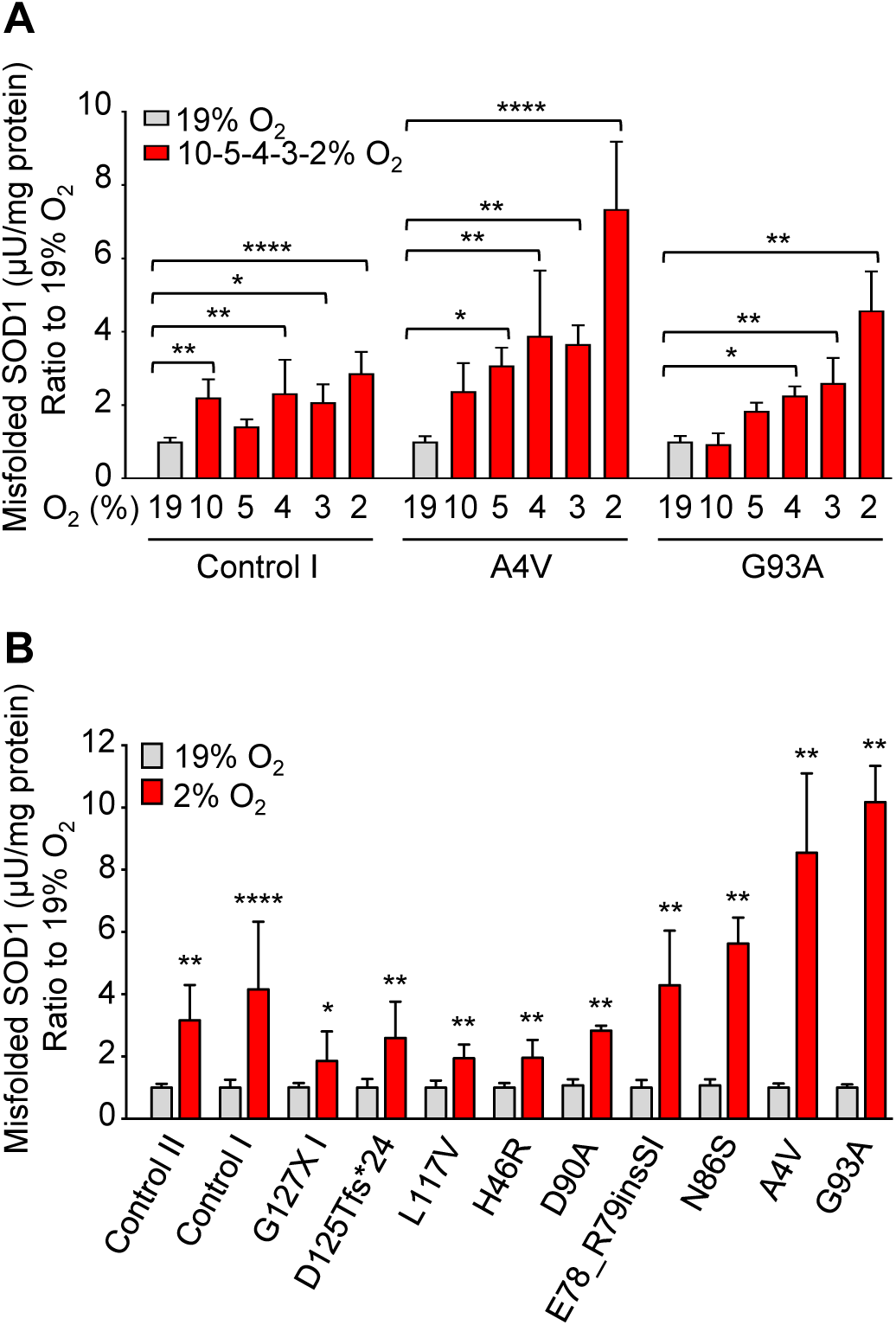
Enhanced misfolding response in patient-derived MNACs. Misfolded SOD1 quantified in MNAC extracts by misELISA, normalized to total protein and presented as a ratio to the level present in replicate cultures maintained at 19% O_2_ for 24 h. (A) O_2_ tension (19-2%) dependent increases in misfolded SOD1. Data are expressed as the mean ± SD (n = 6; 2 replicate experiments from independent differentiations, each with 3 replicate cultures), \**p* < 0.05, \*\**p* < 0.01, \*\*\*\**p* < 0.0001, analyzed by Kruskal-Wallis test followed by Dunn’s test. (B) SOD1 misfolding is increased in MNACs following exposure to 2% O_2_ for 24 h. Data are expressed as the mean ± SD (n = 6 to 18; 2-6 replicate experiments from independent differentiations, each with 3 replicate cultures), \**p* < 0.05, \*\**p* < 0.01, \*\*\*\**p* < 0.0001, analyzed by Mann-Whitney U test.

Large increases in SOD1 misfolding, ranging up to 10-fold, were induced in MNACs following exposure to 2% O_2_ for 24 h (Fig 3B and Fig EV4). The largest effects were found in several of the lines expressing full-length mutant SOD1s (N86S, A4V, and G93A). However, in contrast to fibroblasts a robust 3 – 4-fold increase was also seen in controls expressing SOD1^WT^ and in MNACs that were heterozygous for *SOD1^G127X^* or *SOD1^D125Tfs*24^* mutations. This is likely to represent enhanced misfolding of SOD1^WT^ in MNACs (Forsgren et al., under revision), since the level of mutant protein did not increase in the SOD1^G127X^ line (Fig EV5).

Our previous study of patient-derived fibroblasts showed that misfolded SOD1 is primarily degraded by the proteasome (Keskin et al., 2016). However, exposure to low O_2_ tension did not lead to a reduction in proteasome activity in MNAC extracts (Fig EV3C). Nor were there increases in SOD1^G127X^ protein in either soluble (Fig EV5) or detergent-insoluble (Fig EV6B) fractions, which both increased greatly upon proteasome inhibition (Keskin et al., 2016). Inhibition of the proteasome also leads to copious aggregation of SOD1^D125Tfs*24^ in fibroblasts (Keskin et al., 2016), whereas no changes were found here in response to low O_2_ tension (see below; Fig 7). Hence, increased SOD1 misfolding at low O_2_ tension could not be attributed to a reduction in proteasome activity.

### Low oxygen tension promotes reductive cleavage of the disulfide bond

We have shown that reduction of the C57-C146 bond is a prerequisite for SOD1 misfolding in the CNS (Zetterström et al., 2013, Zetterstrom et al., 2007). To examine the status of the disulfide bond in MNACs we determined the proportions of disulfide-reduced and oxidized SOD1 by non-reduced western blotting (Fig 4A and B). Culture at 2% O_2_ for 24 h resulted in significant increases in disulfide-reduced SOD1, both in control lines expressing SOD1^WT^ and those expressing mutant SOD1s. However, this was not the case for the SOD1^H46R^ line, where a large proportion of the mutant protein is known to be disulfide-reduced under control conditions (Winkler et al., 2009).

**Figure 4.**
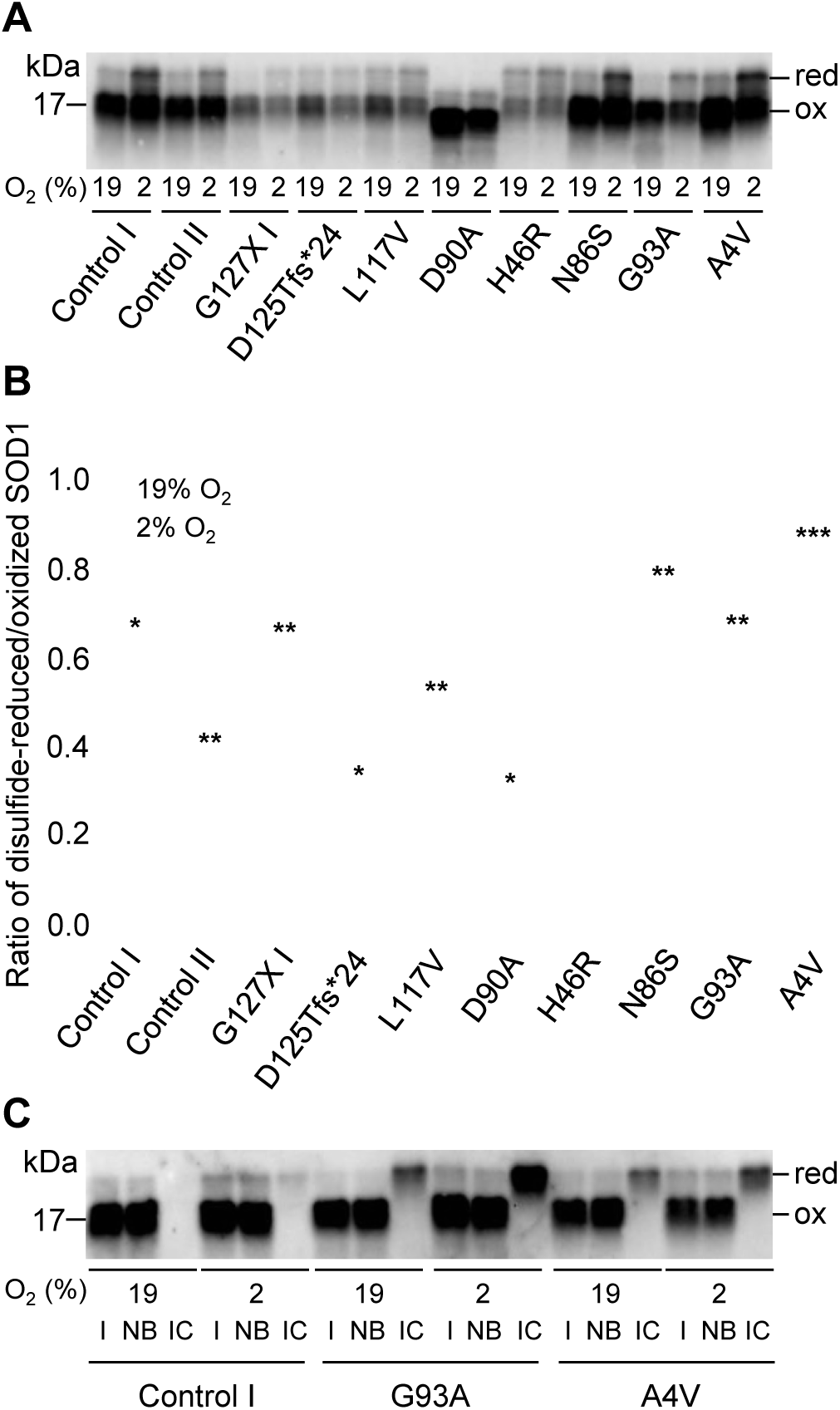
Low O_2_ tension promotes disulfide bond reduction and misfolding of SOD1. Analysis of MNAC extracts following exposure to 19% or 2% O_2_ for 24 h. (A) Disulfide-reduced (red) and -oxidized (ox) SOD1 were resolved by non-reducing SDS-PAGE followed by western blotting with an anti-SOD1 24-39 aa antibody. (B) Scatter plot showing the ratios of disulfide-reduced to -oxidized SOD1 at 19% (grey circles) O_2_ versus 2% (red circles). Data are expressed as the mean ± SD (n = 4 to 8; 2-6 replicate experiments from independent differentiations, each with 3 replicate cultures), \**p* < 0.05, \*\**p* < 0.01, \*\*\**p* < 0.001, analyzed by the Mann-Whitney U test to compare to the respective line cultured at 19% O_2_ for 24 h. (C) Western blot showing analysis of immunocaptured misfolded SOD1. Input (I, 1/40^th^ of the sample), non-bound (NB, 1/40^th^) and immunocaptured (IC, entire sample) fractions of SOD1 from MNACs cultured at 19% O_2_ or 2% O_2_ for 24 h using anti-SOD1 24-39 aa. Disulfide-reduced and -oxidized SOD1 were quantified using anti-SOD1 aa 57-72. Only disulfide-reduced SOD1 was captured.

### Misfolded SOD1 in MNACs lacks the disulfide bond

To confirm that misfolded SOD1 lacks the C57-C146 disulfide bond, we immunocaptured SOD1 from extracts of MNACs, which had been cultured at 19% or 2% O_2_, using the same antibody (SOD1 aa 24-39) and capture conditions (1 h at 23°C) used in the misELISA. This antibody reacts with disordered SOD1, independent of the C57-C146 disulfide status (Forsberg et al., 2010). Disulfide-reduced and oxidized SOD1 were quantified by non-reduced western blotting (Fig 4C). Input samples from control, SOD1^G93A^ and SOD1^A4V^ MNACs contained a majority of oxidized and a small proportion of reduced SOD1, which increased at low O_2_ conditions. Notably, the misfolded SOD1-specific antibody only captured disulfide-reduced SOD1. In control MNACs, this was not detectable at 19% O_2_ but increased at 2% O_2_ to ~1.4% of the disulfide reduced SOD1 in the input sample. In SOD1^G93A^ and SOD1^A4V^ MNACs, immunocaptured disulfide reduced misfolded SOD1 increased substantially (to 21% and 10% of the input samples at 2% O_2_, respectively). Hence, in patient-derived MNACs, the majority of disulfide-reduced SOD1 retained an ordered structure. This agrees with observations in the spinal cord of SOD1 Tg mouse models, where approximately 5 – 10% of disulfide-reduced human SOD1 is disordered (Zetterström et al., 2013, Zetterstrom et al., 2007).

### Low oxygen tension does not alter determinants of SOD1 redox status

Oxidation of the C57-C146 disulfide bond is catalyzed by CCS (Culotta et al., 1997, Fetherolf et al., 2017, Furukawa et al., 2004), and it can be reduced by a mechanism involving reducing equivalents from GSH via glutaredoxin-1 (Carroll et al., 2006). To determine whether these factors were affected by low O_2_ tension and responsible for increased misfolding, we analyzed their levels in MNACs grown at 19% vs 2% O_2_. No significant changes were detected in CCS or glutaredoxin-1 levels by western blotting (Fig 5A and B and Fig EV7). We next quantified GSH and GSSG in MNACs and fibroblasts (Fig 5C and *D*). GSH and GSSG concentrations were higher in MNACs than in fibroblasts, but exposure to low O_2_ tension did not lead to significant changes in GSH or GSSG levels. The concentrations of GSSG were remarkably high in MNACs, resulting in very low GSH/GSSG ratios, but these were not affected by O_2_ (Fig 5E). Hence, low O_2_ tensions do not induce disulfide reduction and misfolding of SOD1 via gross perturbations of the GSH/GSSG redox couple (Schwarzlander et al., 2016).

**Figure 5.**
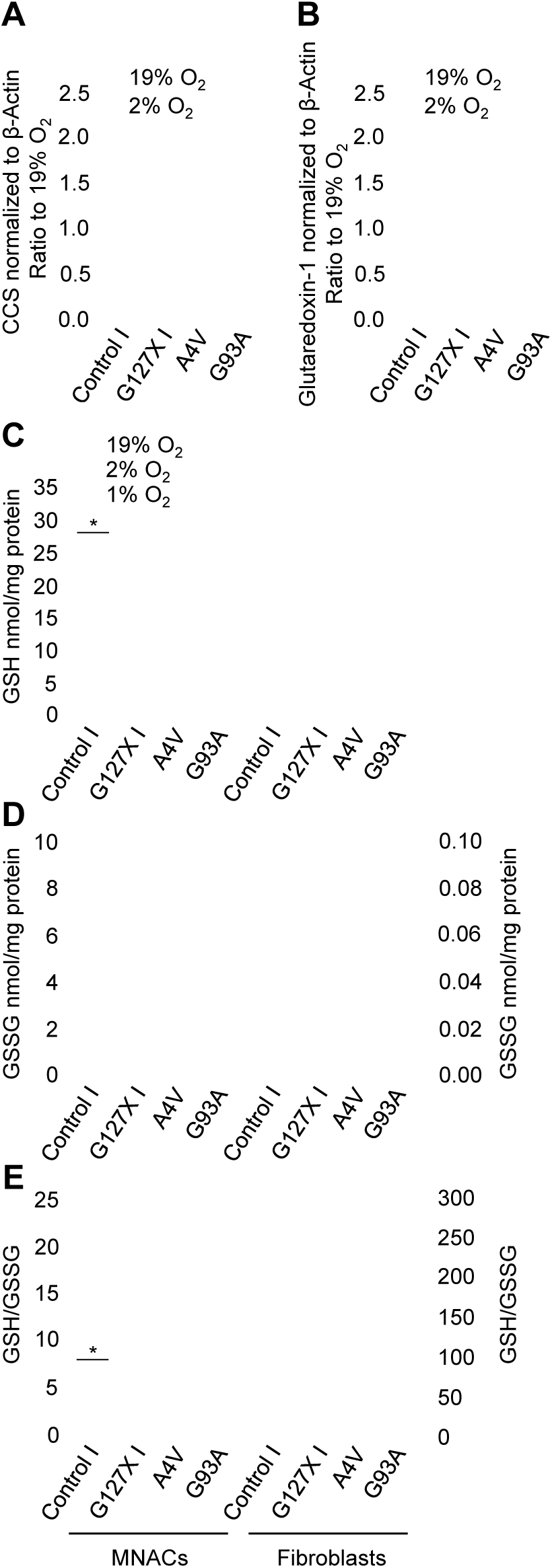
Low O_2_ tension does not alter known determinants of SOD1 C57-C146 redox status. (A) CCS and (B) glutaredoxin-1 quantified in MNAC extracts by western blotting and normalized to β-actin as a loading control. Cells exposed to 19% O_2_ (grey bars) or 2% O_2_ (red bars) for 24 h. Plotted as a ratio of the level present in replicate cultures maintained at 19% O_2_ for 24 h. Data are expressed as the mean ± SD (n = 4 to 10; 2-6 replicate experiments from independent differentiations, each with 3 replicate cultures), analyzed by Mann-Whitney U test. (C) GSH levels, (D) GSSG levels and (E) GSH/GSSG ratios determined in MNACs (left, red bars) and fibroblasts (right, blue bars). Data are expressed as the mean ± SD (n = 6; 2 replicate experiments from independent differentiations, each with 3 replicates for MNACs) and (n = 3 replicate cultures of fibroblasts), \**p* < 0.05, analyzed by the Mann-Whitney U test to compare to the respective line cultured at 19% O_2_ for 24 h.

### Maintenance of the disulfide bond and SOD1 structure is oxygen-dependent

Increased misfolding at low O_2_ tension could be due to impaired disulfide oxidation of newly synthesized SOD1, or to reduction of the disulfide bond in the mature protein. To distinguish the relative contribution of these two possible mechanisms, we compared the misfolding response in fibroblasts cultured in the absence or presence of the protein synthesis inhibitor cycloheximide (CHX) for 24 h. No significant effect of CHX on the level of misfolded SOD1^WT^ was seen in control fibroblasts cultured at 19% or 1% O_2_ (Fig 6A). In contrast, CHX treatment led to a large reduction in misfolded SOD1 in the SOD1^G127X^ line at both O_2_ tensions (Fig 6B). This confirmed that both the inhibition of protein synthesis, and the degradation of the disordered SOD1^G127X^ mutant protein, was efficient. In the SOD1^G93A^ line, CHX treatment resulted in a significant reduction in misfolded SOD1 at both O_2_ tensions. Hence, a proportion of misfolded SOD1 resulted from a lack of disulfide oxidation in newly synthesized SOD1^G93A^ (Fig 6C). However, the misfolding response was enhanced at 1% compared to 19% O_2_ even when protein synthesis was inhibited. Thus, O_2_ is also required for maintenance of the disulfide bond and structure of the existing pool of mature SOD1.

**Figure 6.**
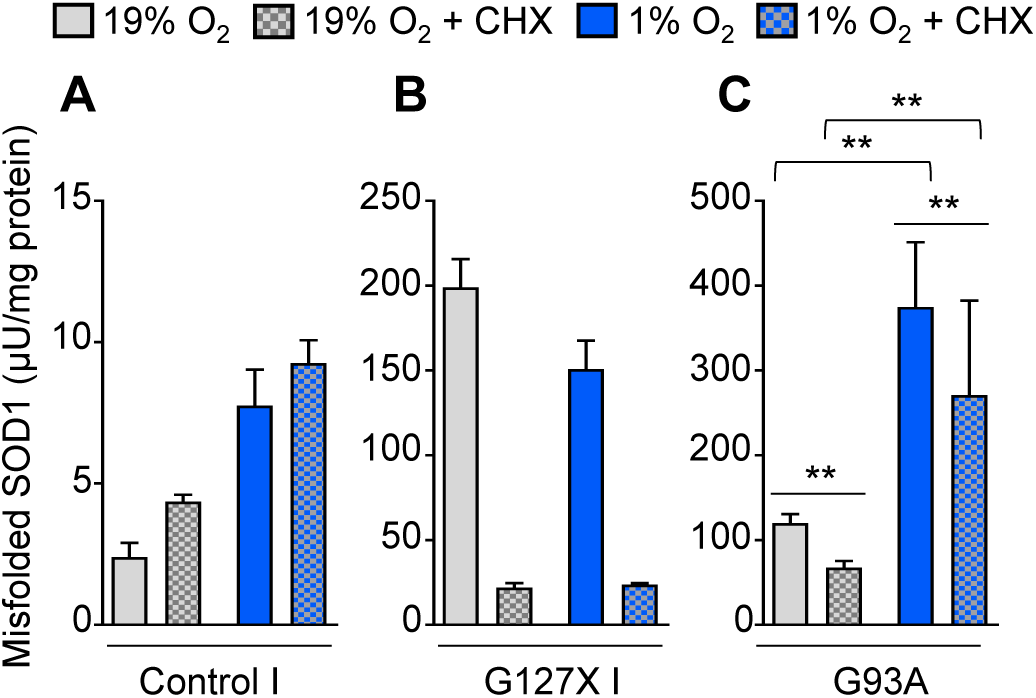
Low O_2_ tension increases disulfide bond reduction in both nascent and mature SOD1. Misfolded SOD1 quantified in; (A) Control I, (B) SOD1^G127X^ and (C) SOD1^G93A^ fibroblast extracts by misELISA, normalized to total protein. Cells grown at 19% O_2_ (grey bars) or 1% O_2_ (blue bars) for 24 h in the absence, or presence (checker board pattern), of cycloheximide (CHX; 50 µg/ml). Data are expressed as the mean ± SD (n = 3 replicate cultures for Control I and SOD1^G127X^) and (n = 6; 2 replicate experiments each with 3 replicate cultures for SOD1^G93A^ normalized by division by the means of each experiment), \*\**p* < 0.01, analyzed by the Mann-Whitney U-test.

### Low oxygen tension promotes SOD1 aggregation

Since disulfide reduction and misfolding promote SOD1 aggregation (Chattopadhyay et al., 2008, Furukawa et al., 2008, Lang et al., 2012), we quantified detergent-resistant SOD1 aggregates in both fibroblasts and MNACs. Low levels of aggregation were found in fibroblasts, which did not increase significantly in response to low O_2_ (Fig 7). In contrast, we found that low O_2_ tension induced marked increases in aggregation in MNACs carrying full-length mutant SOD1s (E78_R79insSI, N86S, G93A and A4V), but not in SOD1^WT^ control lines or SOD1^L117V^, which has SOD1^WT^-like stability (Synofzik et al., 2012). The *SOD1^H46R^* mutation disrupts copper binding catalyzed via CCS and impairs formation of the disulfide bond (Winkler et al., 2009). In agreement, the high level of aggregation present at 19% O_2_ in the SOD1^H46R^ line did not change at 2% O_2_. Neither of the lines expressing the C-terminally truncated mutants that lack the disulfide bond (SOD1^G127X^ and SOD1^D125Tfs*24^), showed increased aggregation. Hence, increased aggregation correlated closely with increased misfolding in MNACs expressing full-length mutant SOD1s.

**Figure 7.**
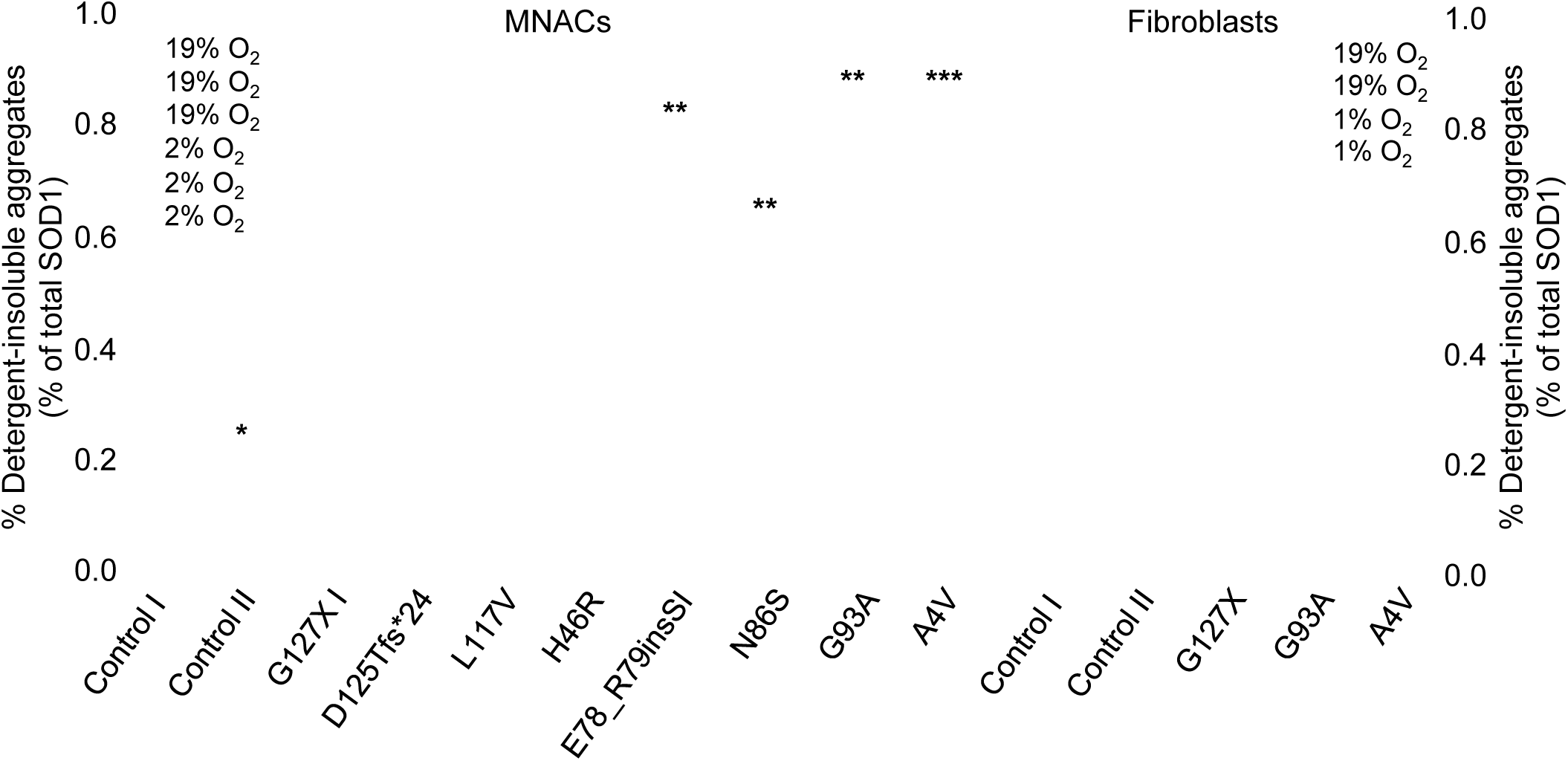
Low O_2_ tension promotes SOD1 aggregation in MNACs but not in fibroblasts. Scatter plots show the amounts of SOD1 present in the detergent-insoluble fraction determined by western blotting as a % of soluble SOD1 present in the cell extract determined by ELISA; MNACs (red; 2% O_2_/24 h) and fibroblasts (blue; 1% O_2_/24 h) in comparison to replicate cultures (grey; 19% O_2_/24 h). Full-length SOD1s are depicted as filled circles. The aa 57-72 antibody used for detection showed relatively strong bands in the SOD1^D125Tfs*24^ samples (Fig EV6A). This represents mutant SOD1 (filled squares), since negligible bands were detected using an antibody raised against a C-terminal SOD1 (aa 143-153) peptide (Fig EV6E). The SOD1^G127X^ mutant protein (filled triangles) was quantified using a hSOD1^G127X^-specific antibody (Fig EV6 B and D). Data are expressed as the means ± SD (n = 4 to 10; 2-6 replicate experiments from independent differentiations, each with 3 replicates for MNACs) and (n = 3 to 6; 1-3 experiments each with 3 replicates for fibroblasts), \**p* < 0.05, \*\**p* < 0.01, \*\*\**p* < 0.001, analyzed by the Mann-Whitney U test.

## Discussion

The principal finding of this study is that culture at low O_2_ tension leads to remarkably large increases in SOD1 C57-C146 disulfide bond reduction and misfolding. In addition to being required for bond formation, we have identified that O_2_ is also critical for its maintenance. Low O_2_ tension resulted further in enhanced aggregation in MNACs but not in fibroblasts. This supports the idea that in human patient-derived models, as well as in Tg mice, the C57-C146 disulfide bond is indeed an ALS-related Achilles heel of SOD1 in the reducing environment of the cytosol.

The O_2_ tensions in the gas phase found to enhance SOD1 misfolding are within the range considered to be normoxic in human and animal tissues. However, the range of O_2_ tensions present within cultured cells are likely to be lower. This is due to both the rate of O_2_ diffusion through the culture medium and the rate of O_2_ consumption by cellular respiration, which depends on both cellular mass and the rate of oxidative metabolism (Pettersen et al., 2005). Because these variables are difficult to control for between different cell lines, we chose to compare replicate samples cultured at different O_2_ tensions. The cells, which had been propagated and adapted to growth at 19% O_2_, were tested within a range that was found not to cause overt toxicity.

A study in young resting awake mice has indicated that O_2_ tensions in the CNS vary between 0.2 – 5% (Lyons et al., 2016). Even lower levels were recorded under normal conditions, e.g. in periarteriolar areas. Considering this great variation found in young mice, it is reasonable to assume that localized hypoxia could occur transiently, or more chronically, in elderly humans even in the absence of overt disease.

Apart from the strong risk associated with aging, other suggested non-hereditary risk factors for ALS include smoking (Gallo et al., 2009), CNS trauma, particularly to the motor cortex (Rosenbohm et al., 2014), embolisations of arteriovenous malformations, transient ischaemic attack, and stroke (Turner et al., 2016). Strenuous physical activity has also been suggested to increase the risk for ALS (Chio et al., 2005), although this is contentious (Gallo et al., 2016). The mechanisms underlying these risk factors are not understood, but a unifying characteristic could be transient or chronic CNS hypoxia due to reduced vascular perfusion.

In *SOD1* ALS mouse models, the average lifespan of homozygous Tg mice is approximately half that of hemizygous animals (Jaarsma et al., 2000, Jonsson et al., 2004). A dose-dependence on mutant SOD1s also exists for the development of ALS in humans. Individuals that are homozygous for the *SOD1* mutations L84F, N86S or L126S show earlier onset and more rapid progression than those that are heterozygous for the same mutation (Alavi et al., 2013, Boukaftane et al., 1998, Hayward et al., 1998, Kato et al., 2001). Furthermore, the D90A mutant protein, which has wild type-like stability, typically only causes ALS in homozygous individuals (Andersen et al., 1996). Hence, a clear link exists between a doubling of the load of mutant SOD1 and disease initiation, or progression. The large increases in misfolded SOD1 in response to low O_2_ tension we report here, particularly in MNACs, could contribute to the initiation and progression of ALS.

Except for SOD1^D90A^, the cell lines tested here were derived from individuals heterozygous for *SOD1* mutations. Whereas SOD1^D90A^ has a larger electrophoretic mobility, the other full-length SOD1 mutants show the same mobility as wild-type SOD1. The SOD1^D125Tfs*24^ mutant protein also has the same mobility as SOD1^WT^ (Keskin et al., 2016). Therefore, these SOD1 mutants cannot be distinguished from SOD1^WT^ by western blotting or in the misELISA (Keskin et al., 2016). Owing to widely different stabilities and rates of degradation, the levels of mutant SOD1s in cultured human cells vary from comparable to the wild type protein, to very low (Keskin et al., 2016). Whereas there were no large differences in the proportion of disulfide-reduced SOD1 between *SOD1* mutant and control MNACs, both basal and low O_2_-enhanced levels of misfolded and aggregated SOD1 were much greater in the *SOD1* mutant cultures. For the most part, mutant SOD1s most likely contributed substantially to these enhanced levels.

Recent findings *in vivo* suggest that prion-like growth and spread of SOD1 aggregation could be the primary disease mechanism of SOD1-induced ALS (Ayers et al., 2014, Ayers et al., 2016, Bidhendi et al., 2016). Disordered SOD1 species would be critical substrates for both the nucleation and growth of prion-like aggregates. This would have a greater probability of occurring in areas of the CNS with sustained low O_2_ tension. Consistent with this, even a short (24 h) exposure to low O_2_ tension resulted in large increases in SOD1 aggregation in patient-derived MNACs, which do not overexpress the protein. Typically SOD1 overexpression is required for substantial aggregation to occur (Prudencio et al., 2009).

Significant increases in disulfide-reduced and misfolded SOD1 were also seen in cells from healthy control individuals and cells derived from patients carrying ALS-linked mutations in other genes. This could be important since there is mounting evidence for involvement of SOD1^WT^ in ALS patients without *SOD1* mutations. Although this is debated (Da Cruz et al., 2017), several studies have demonstrated inclusions of misfolded SOD1^WT^ in the soma of motor neurons from sporadic as well as familial ALS patients carrying ALS-linked mutations in other genes (Bosco et al., 2010, Forsberg et al., 2011, Forsberg et al., 2010, Pokrishevsky et al., 2012). Furthermore, co-culture of motor neurons with primary astrocytes derived from sporadic ALS patients indicates an involvement of SOD1^WT^ in the disease (Haidet-Phillips et al., 2011), and mice that express human SOD1^WT^ at high rate develop both SOD1 aggregation and a fatal ALS-like disease (Graffmo et al., 2013).

SOD1s carrying mutations that affect the C57-C146 disulfide bond should not be affected directly by low O_2_ tension. However, in several mutant *SOD1* Tg models coexpression of human SOD1^WT^ has been found to exacerbate the disease (Deng et al., 2006, Jaarsma et al., 2000, Prudencio et al., 2010, Wang et al., 2009). Low O_2_ tension effects on SOD^WT^ might contribute to the pathology in heterozygous individuals, perhaps through coaggregation of wild-type and mutant SOD1s.

In conclusion, we show that O_2_ tension is a principal determinant of SOD1 misfolding and aggregation. Our findings suggest that CNS areas with low O_2_ tension could act as foci for the initiation or progression of ALS. This mechanism might contribute to the enhanced risk for the disease associated with aging, as well as other factors that impair vascular perfusion.

## Materials and Methods

### Human materials

Samples from patients and non-disease controls (Table EV1), including blood samples for genotyping and skin biopsies for fibroblast culture, were collected with approval of the Regional Ethical Review Board in Umeå and in accordance with the principles of the Declaration of Helsinki (WMA, 1964), following written informed consent.

### Reagents and chemicals

All reagents and chemicals were obtained from Sigma or Thermo Fisher Scientific unless stated otherwise.

*SOD1*, *FUS*, *TBK1* and *C9orf72* genotyping

*SOD1* and *C9orf72* genotyping were done as described (Keskin et al., 2016). For *FUS*, only exons 2-6 and 11-15 were analyzed. *TBK1* genotyping was performed as described (Freischmidt et al., 2015). All individuals tested negative for mutations in a panel of other ALS-linked genes (details available upon request).

### Derivation of human fibroblasts

Following screening of blood to identify *SOD1, FUS, TBK1* and *C9orf72* mutation carriers, fibroblasts were generated from a 3 mm punch skin biopsy (upper arm) from 10 ALS patients with *SOD1* mutations (A4V, H46R, E78_R79insSI, N86S, D90A, G93A, L117V, D125Tfs*24 and G127Gfs*7 (G127X – in 2 patients)). One ALS patient with Q23L mutation in *FUS*, 5 ALS patients with *TBK1* mutations (A417X – in 2 patients, M598V, I450Kfs*14, p.690-713del), 1 ALS and 1 FTD patient with massive intronic GGGGCC repeat-expansions in *C9orf72*, and 4 non-disease control individuals (Table EV1). The establishment of the lines followed standard procedures (Keskin et al., 2016). ALS patients were diagnosed according to EFNS guidelines (Andersen et al., 2005). All patients were heterozygous for their corresponding mutations except the *SOD1^D90A^* patient, who was homozygous. Control subjects were relatives of ALS patients. All healthy control subjects tested negative (wt/wt) for a panel of ALS-associated genes including *SOD1, C9orf72, TBK1* and *UBQLN2*.

### Generation and maintenance of iPSCs

Fibroblast lines were reprogrammed either with the mRNA Reprogramming Kit (Stemgent, Cambridge, MA, USA) (Warren et al., 2010) using a commercial service (Cellectis AB, Gothenburg, Sweden) or using an episomal vector system (Okita et al., 2011) (Table EV1). IPSCs were cultured using the DEF-CS culture system (Takara Bio Europe, Gothenburg, Sweden) seeded at a density of 40,000 cells/cm^2^. Media changes were preformed every 24 h and cells were passaged after 3-4 days at a density of 1.5 -2x10^5^ cells/cm^2^ using TrypLE and plated in DEF media supplemented with GF1, GF2 and GF3 (Takara Bio Europe, Gothenburg, Sweden).

### Generation of iPSC-MNACs

To generate MNACs, iPSCs at 90% confluence were maintained with N2/B27 media, consisting of DMEM/F12, Neurobasal, 1x Nonessential Amino Acids (NEAA; Millipore, Bedford, MA, USA), 2 mM L-glutamine, 1% (v/v) N2 supplement, 2% (v/v) B27 supplement and penicillin/streptomycin. Over 14 days of differentiation, N2/B27 media were supplemented with 1 μM all-*trans* retinoic acid (RA) and 1 μM smoothened agonist (SAG; Millipore, Bedford, MA, USA). For the first 6 days, cells were kept in N2/B27 media, supplemented with 10 μM SB431542 and 100 nM LDN (Stemgent, Lexington, MA, USA). Dual SMAD inhibition (SB and LDN) was maintained until day 7, after which cells were maintained with N2/B27, supplemented with 4 μM SU5402 (Stemgent, Lexington, MA, USA) and 5 μM DAPT (Selleck Chemicals, Houston, TX, USA) (Lamas et al., 2014).

At Day 14, differentiated cells (neurons and astrocytes) were dissociated to single cells with Accutase and plated onto poly-L-ornithine/laminin-coated 6-well plates (BD Biosciences, Franklin Lakes, NJ, USA) at a density of 100,000 cells/cm^2^ or 13 mm diameter coverslips (No. 1.5, VWR, Stockholm, Sweden) at 40,000 cells/cm^2^. MNAC cultures were maintained in MN culture media, consisting of Neurobasal, NEAA, 2 mM L-glutamine, 1% (v/v) N2 supplement, 2% (v/v) B27 supplement, 0.4 mg/L ascorbic acid 25 µM glutamate, 25 µM 2-mercaptoethanol, 1 μM RA, 20 µM Y-27632 (Abcam, Cambridge, UK) and penicillin/streptomycin. Cells were incubated in a humidified atmosphere at 37°C supplemented with 5% (v/v) CO_2_ and MNAC cultures. On days 11 -13 days after plating (days 24-26 of the differentiation), MNACs were exposed to different O_2_ tensions prior to analysis.

### Exposure to different oxygen tensions

Fibroblasts were plated at 8000 cells/cm^2^ in 60 mm dishes or 6-well plates (BD Biosciences, Franklin Lakes, NJ, USA) and incubated in an humidified atmosphere at 37°C with 5% (v/v) CO_2_. After reaching 60-70% confluence, the medium was replaced prior to exposure to different O_2_ tensions. Differentiated MNACs were exposed to different O_2_ concentrations without medium change. For O_2_ tensions lower than 19%, cells were incubated in a sealed chamber (Biospherix, Lacona, NY, USA) gassed with humidified gas mixtures (1 -10% O_2_, 5% CO_2_, 94 -85% N_2_) at 37°C. The O_2_ concentration was maintained using a ProOx P110 oxygen controller (Biospherix, Lacona, NY, USA) supplied with N_2_ gas. For each experiment cells were also cultured in parallel under humidified atmospheric O_2_ (19% O_2_, 5% CO_2_, 76% N_2_) in a standard tissue culture incubator.

### Generation of patient-derived primary astrocytes

A piece (2 cm) of ventral horn from the thoracic spinal cord was dissected for cell isolation as previously described (Haidet-Phillips et al., 2011). The tissue was diced and dissociated using 2.5 U/ml papain and 0.5 U/ml Dispase (Stemcell Technology, Canada, Inc.) in 1xHBSS supplemented with 40 μg/ml DNAse at 37°C for 45 min with mixing every 5 min Cells were dissociated using a P1000 pipette tip and then mixed with KnockOut DMEM/F12 media containing 10% (v/v) foetal bovine serum (FBS), before filtering the cells through a 0.7 µm cell strainer followed by centrifugation. The cell pellet was resuspended in DMEM/F12 + 10% FBS and mixed with an equal volume of Percoll (GE healthcare, Chicago, Illinois). The mixture was centrifuged at 20,000 *g* for 30 min at 4°C and the flocculent layer above the red blood cell layer was collected, washed, resuspended and cultured in primary cell media containing KnockOut DMEM/F12 + 10 % (v/v) FBS supplemented with 20 ng/ml FGF-2, 20 ng/ml EGF, 20 ng/ml PDGF-AB (all from Peprotech, Rocky Hill, NJ), 2 mM GlutaMAX Supplement, 1x (v/v) StemPro Neural Supplement and penicillin/streptomycin. The cells were cultured on CELLstart-coated plates and after 24 hours the medium was replaced with serum-free primary cell media. Every 2 days, half of the medium was replaced until cells reached 80% confluence after 4-5 weeks. The cells were then passaged, expanded and plated either in coated 6-well plates (BD Biosciences, Franklin Lakes, NJ, USA), for exposure to different O_2_ tensions, or 13 mm coated coverslips (VWR, Stockholm, Sweden) for immunohistochemistry.

### Cell collection and detergent-insoluble aggregates

Cells were washed at room temperature with PBS containing 40 mM iodoacetamide (IAM), which blocks reduced cysteine residues via alkylation and prevents artificial formation of the C57-C146 disulfide bond (Zetterstrom et al., 2007). The cells were detached with trypsin, and then washed with PBS containing 40 mM IAM and 0.5% (v/v) FBS. After centrifugation of the suspension at 500 *g* for 5 min, the cell pellet was collected and snap frozen on dry ice and stored in a -80°C freezer.

Cell pellets were lysed in ice-cold PBS containing the Complete EDTA-free protease inhibitor cocktail (Roche Diagnostics, Mannheim, Germany), 40 mM IAM and 0.5% (v/v) NP-40 using a Sonifier Cell Disrupter (Branson, Danbury, CT, USA). Lysates were centrifuged at 20,000 *g* for 30 min at 4°C and supernatants were collected and analyzed directly with by misELISA (see below). Pellets were washed twice in ice-cold lysis buffer containing 0.5% (v/v) NP-40 to remove remaining detergent-soluble proteins. The final pellets were used for determination of detergent-insoluble SOD1 aggregates by western blotting.

### Immunocapture

A rabbit anti-human SOD1 antibody raised against aa 24-39 of SOD1 was cross-linked to Dynabeads^®^ M-270 Epoxy with the Dynabeads Antibody Coupling Kit (Invitrogen). Beads were recovered with a magnet, washed with the supplied buffers to remove unbound antibody and equilibrated with PBS containing the Complete EDTA-free protease inhibitor cocktail (Roche Diagnostics, Mannheim, Germany), 40 mM IAM and 0.5% (v/v) NP-40. Antibody-coated beads were incubated with 20,000 *g* cell lysate supernatants for 1 h at 23°C. Beads were washed five times with PBS containing the Complete EDTA-free protease inhibitor cocktail (Roche Diagnostics, Mannheim, Germany), 40 mM IAM and 0.5% (v/v) NP-40 and samples were eluted by boiling in 1x sample buffer containing 40 mM IAM. A proportion of the input and non-bound fractions (1/40^th^), as well as the entire immunocaptured fractions, were analyzed using non-reduced western blotting to determine the proportions of reduced and oxidized SOD1.

### Analysis of misfolded and total SOD1 by ELISA

Misfolded SOD1 ELISA (misELISA) was carried out as described (Zetterström et al., 2011, Zetterström et al., 2013). A primary rabbit antibody was raised against a peptide corresponding to aa 24-39 of the human SOD1 sequence. This antibody reacts only with disordered SOD1 species and lacks affinity for the natively folded protein (Bergh et al., 2015, Forsberg et al., 2011, Forsberg et al., 2010). A goat anti-human SOD1 antibody was used as a secondary antibody. It was raised against SOD1 that had been denatured by incubation with guanidinium chloride and EDTA, and reacts preferentially with the disordered protein (Zetterström et al., 2011). For calibration of the misELISA, a fresh spinal cord from a transgenic mouse expressing hSOD1^G127X^ was homogenized in 25 volumes 10 mM K phosphate, pH 7.0 in 0.15 M NaCl, containing the Complete protease inhibitor cocktail (Roche Diagnostics, Mannheim, Germany) and 40 mM IAM. After centrifugation at 20,000 *g*, the supernatant was divided into multiple aliquots that were stored at -80°C. One unit of misfolded SOD1 is defined as the amount present in 1 g wet weight of the original human SOD1^G127X^ standard spinal cord.

The misELISA only reacts with disordered/misfolded SOD1 species. There is no response to holoSOD1 or SOD1s that lack Cu and/or the C57-C146 disulfide bond as long as the polypeptide is natively folded. The cell extracts were incubated for 1 h at 23°C with the capture antibody in the misELISAs. This temperature was selected because at 37°C, some folded SOD1 species have been found to unfold continuously in diluted extracts, such as the cell extracts analyzed here (Zetterström et al., 2013). Moreover, it has been shown that mutations can influence the conformations of immature SOD1 states which could affect the reactivity with the antibodies (Santamaria et al., 2016, Sekhar et al., 2016). Thus, the actual levels of misfolded SOD1 at physiological temperature are not mirrored equally for the SOD1 variants, and misELISA results are not directly comparable between cell lines expressing different SOD1s. However, for a given cell line expressing the same SOD1 variant, the effects of different O_2_ tensions on the levels of misfolded SOD1 can be determined.

Total SOD1 was quantified in extracts with a sandwich ELISA based on rabbit capture and goat detection antibodies, both raised against native human SOD1 (Zetterström et al., 2011). It was standardized against human hemolysate with a known SOD1 content calibrated against pure human SOD1, the concentration of which was determined by quantitative amino acid analysis (Marklund et al., 1997).

### Western blot, quantification of SOD1 protein, and determination of disulfide-reduced SOD1

The total protein content of cell lysates was determined using the BCA Protein Assay Kit. Western blots were performed in Any kD and 18% (w/v) Criterion TGX precast gels (BioRad Laboratories, Hercules, CA, USA) as previously described (Keskin et al., 2016).

For analysis of SOD1, the primary antibodies used were: rabbit anti-human SOD1 antibodies raised against peptides corresponding to aa 24-39 (2.3 μg/ml), 57–72 (1.6 μg/ml), aa 144-153 (5.2 μg/ml), aa 123-127GQRWK (4.8 µg/ml, G127X-specific) (Jonsson et al., 2006). The same human SOD1 standard described above was used for calibration. For analysis of SOD1^G127X^ using the aa 123-127GQRWK antibody, blots were standardized with an hSOD1^G127X^ transgenic mouse extract in which the SOD1^G127X^ content was determined by western blot using the human-specific aa 24-39 antibody. The proportions of disulfide-reduced and oxidized SOD1 were determined using non-reducing western blotting, i.e. omitting reductant and adding 40 mM IAM to the sample buffer as described (Jonsson et al., 2006, Zetterström et al., 2013, Zetterstrom et al., 2007).

Other antibodies used were rabbit anti-CCS raised against peptides corresponding to aa 252–270 of the human CCS sequence (CCS, 0.9 μg/ml) (Jonsson et al., 2006), rabbit anti-CCS (1:1000 Santa Cruz Biotechnology, Dallas, TX, USA), rabbit anti-glutaredoxin-1 (1:250, Abcam, Cambridge, UK), mouse anti-β-actin (1:200,000; Millipore, Bedford, MA, USA) and rabbit anti-GAPDH (1:1000, Abcam, Cambridge, UK). Secondary antibodies used were horseradish peroxidase (HRP)-conjugated polyclonal anti-mouse or anti-rabbit IgG (1:10,000, Dako, Glostrup, Denmark). ECL Select reagent (GE Healthcare Biosciences, Piscataway, NJ, USA) was used to detect the signal, which was recorded on a ChemiDoc Touch apparatus (BioRad Laboratories, Hercules, CA, USA) and analyzed using ImageLab software (BioRad Laboratories, Hercules, CA, USA).

### Effect of cycloheximide on SOD1 misfolding

Fibroblasts were exposed to 19% O_2_ or 1% O_2_ as described above in the presence and absence of protein synthesis inhibitor CHX (50 µg/ml) for 24 h prior to harvest.

### Analysis of reduced and oxidized glutathione

After exposure of MNACs and fibroblasts to different O_2_ tensions (1-2-19% O_2_) for 24 h as described above, media was aspirated rapidly and 300 µL of ice-cold extraction mixture (0.25 µM glutathione-(glycine-^13^C_2_,^15^N; GSH-IS), 2.5% metaphosphoric acid, 1 mM EDTA and 0.1% formic acid) were added immediately to the cells. After scraping, the resulting cell suspensions were snap-frozen and stored at -80°C until analysis.

For analysis, a tungsten bead was added to the tubes, which were then homogenized in a bead mill (Retsch MM400) at 30 oscillations/s for 1 min, followed by centrifugation at 22,000 *g* for 20 min Fifty µL of the resulting supernatant was transferred to a LC-MS vial and 1 µL was injected and analyzed by LC-ESI-MSMS (1290 Infinity system from Agilent Technologies, with an Acquity UPLC HSS T3 column, thermostatted to 40°C and coupled to an Agilent 6490 Triple quadrupole mass spectrometer). The analysis was performed as essentially as described (Cao et al., 2013) by comparison to GSH and GSSG solutions made in water (2.5 mM metaphosphoric acid, 1 mM EDTA, 0.1% formic acid) at twelve different calibration concentrations (0.01-10 µM) and containing the GSH-IS.

GSH and GSSG concentrations were normalized to the protein content of the corresponding 22,000 *g* pellets after extraction. Pellets were resuspended by sonication/boiling in sample buffer. Protein estimation was performed using the BCA Protein Assay Kit and bovine serum albumin standards boiled in 1x sample buffer.

### Immunocytochemistry

Cells were fixed in 3.8% (w/v) formaldehyde for 10 min at room temperature, and blocked with 10% (v/v) FBS in PBS containing 0.1% (v/v) Triton X-100 for 1 h at room temperature. Cells were incubated with primary antibodies overnight at 4°C. The primary antibodies used were; anti-neuron-specific class III beta-tubulin (TUBB3, TUJ1, 1:7500, Covance Inc. Princeton, NJ, USA), microtubule-associated protein 2 (MAP2; 1:500, Millipore, Bedford, MA, USA), SMI32 (1:1000, Covance Inc. Princeton, NJ, USA), ISL1/2 (39.4D5,1:5, developed by T.M. Jessell and S. Brenner-Morton, Developmental Studies Hybridoma Bank, created by the NICHD of the NIH and maintained at The University of Iowa, Department of Biology, Iowa City, IA, USA), S100β (1:200) and vimentin (1:1000 Progen, Heidelberg, Germany). Next day, the coverslips were washed and incubated with Alexa-Fluor conjugated secondary antibodies (1:1000) for 1 h at room temperature, and nuclei were counterstained with 4’,6-diamidino-2-phenylindole (DAPI; 0.3 µM). Cells were mounted in Aqua-Polymount (Polysciences, Inc., Warrington, PA, USA).

### In vitro cell cytotoxicity assays

Dead cells that had lost plasma membrane integrity were quantified using the luminogenic substrate AAF-Glo (CytoTox-Glo Cell Viability Assay, Promega, Madison, WI, USA) as previously described (Keskin et al., 2016). Cell viability was calculated by subtracting the luminescence signal obtained before, from the total luminescence values obtained after, permeabilization with digitonin according to the manufacturer’s protocol.

Cellular ATP content was determined using the CellTiter-Glo Luminescent Cell Viability Assay (Promega, Madison, WI, USA).

### Proteasome activity assay

A cell-based luminescent proteasome assay (Proteasome-Glo, Promega, Madison, WI, USA) was used to measure chymotrypsin-like proteasome activity as described (Keskin et al., 2016). The assay contains the luminogenic peptide substrate Suc-LLVY-aminoluciferin for determination of chymotrypsin-like activity of the proteasome.

### Statistical analyses

Statistical analyses were performed using GraphPad Prism version 6.00 (La Jolla, CA, USA). To test for statistical significance between two groups, the Mann-Whitney U test was used. For statistical significance testing between multiple groups, the Kruskal-Wallis test followed by Dunn’s post hoc test was used. The significance level was set to 0.05. All values are given as means ± SD.

## Acknowledgements

We thank the many individuals that generously provided patient or control samples. We thank Anna Wuolikainen for valuable discussions and Agneta Öberg, Eva Jonsson, Eva Bern, and Matthew Marklund for skillful technical assistance. The study was supported by the Swedish Research Council (VRMH 2015-02804), the Knut and Alice Wallenberg Foundation (2012.0091), the Bertil Hållsten Foundation, the Torsten Söderberg Foundation, the Swedish Brain Fund (Hjärnfonden FO2015-0234), the Ulla-Carin Lindquist Foundation, Neuroförbundet, the Stratneuro Initiative, Västerbotten County Council, and the Kempe Foundations. MS was supported by the Else Kröner Fresenius Stiftung.

## Author contributions

Conception and design of the work: IK, EF, ML, UN, PMA, PZ, SLM, JDG. Clinical and pathological diagnoses and sample collection: DJL, MS, TB, PMA. Acquisition, analysis, or interpretation of the data: IK, EF, ML, PMA, TB, UN, PZ, SLM, JDG. Drafting or revising the manuscript for intellectual content: IK, EF, ML, UN, PMA, SLM, JDG with input from all authors. Final approval of the manuscript: All authors.

